# Extruded alginate tubes with myogenic potential

**DOI:** 10.1101/2024.04.30.591971

**Authors:** Cameron M. Lenneman, Emily M. Rose, Brooke A. Strawska, Natalie A. Tyszkiewicz, Karen Dean-Christie, Erin Katz, Joseph M. Roche, Antoine de Morree, Renuka Roche, Mohan E. Tulapurkar, Joseph A. Roche

**Author notes:** **Correspondence to:** Joseph A. Roche, BPT, PhD. Associate Professor. Physical Therapy Program. Department of Health Care Sciences. Eugene Applebaum College of Pharmacy and Health Sciences. Wayne State University. Detroit, MI, USA.

## Abstract

**BACKGROUND:** There are currently no proven methods to reverse muscle loss in humans, which is caused by trauma (e.g., volumetric muscle loss, VML), genetic neuromuscular diseases (e.g., muscular dystrophies, MDs), and accelerated senescence (e.g., sarcopenia). Since muscle tissue is capable of regeneration through muscle satellite cells (MuSCs), the implantation of autologous (or other) donor MuSCs and MuSC-derived myoblasts into host muscles can promote donor-cell-derived myogenesis. Direct injection or implantation of MuSCs or MuSC-derived myoblasts into host muscles only promotes minimal donor-cell-derived myogenesis, whereas implantation of MuSCs/myoblasts along with associated muscle tissue (muscle fibers, extracellular matrix, neurovascular pathways, etc.) gives better results.

**METHODS:** We aim to leverage the benefits of constraining donor myogenic cells within a template that resembles muscle tissue. In this paper, we present a workflow for basic and translational studies aimed at promoting donor-cell-derived myogenesis to increase functional muscle mass in mice. Our workflow involves preparing a slurry of 10% sodium alginate mixed with myogenic cells in cell culture media, extruding the cell-containing slurry into 10% calcium lactate to form tubes, and implanting the cellularized alginate tubes into host muscle.

**RESULTS:** Our data suggest that, the extruded alginate tubes can tolerate a peak stress of 1892 ± 527 mN, that the elastic range is at ~75-125% strain beyond initial length, and that the Young’s modulus (stiffness) is 14.17 ± 1.68 %/mm^2^. Importantly, these mechanical properties render the alginate tubes suitable for a published technique known as minimally-invasive muscle embedding (MIME) that was developed by us to implant myogenic material into host muscle. MIME involves threading donor myogenic tissue into a needle track created within a host muscle. Cellularized alginate tubes implanted into the tibialis anterior muscle of previously euthanized mice had numerous hematoxylin-stained structures similar to nuclear staining, supporting the idea that our alginate tubes can support cell seeding. Alginate tubes that were seeded with MuSCs, incubated in MuSC/myoblast growth (i.e., proliferation) media for two days, incubated in myotube differentiation media for six days, and then minced and reseeded in new dishes, were able to promote in vitro myoblast outgrowth over several days.

**DISCUSSION:** This pilot study is limited in its translational scope because it was performed in vitro and with previously euthanized mice. Additional studies are needed to confirm that cellularized alginate tubes can promote the de novo development of donor-cell-derived muscle fibers, which can contribute to contractile force production.

**CONCLUSION:** Alginate tubes with MuSC/myoblasts can be generated by a simple extrusion method. The alginate tubes have sufficient mechanical strength to tolerate insertion into a host muscle, in a minimally-invasive manner, through a needle track. The cellularized alginate tubes demonstrate myogenic potential since they are capable of being maintained in culture conditions for several days, after which they can still facilitate myoblast outgrowth in a dish.

## BACKGROUND

Regenerative muscle biologists have developed effective methods to grow myogenic cells (i.e., muscle satellite cells and myoblasts) in petri dishes and bioreactors [reviewed in (Roche 2022, Grounds 2014, Smith et al. 2022)]. However, minimal progress has been made in regrowing functional skeletal muscle tissue in living hosts from donor myogenic cells (Mendell et al. 1995, Skuk et al. 2006). The lack of substantial progress in this area limits the chances of individuals with muscle loss from trauma (e.g., volumetric muscle loss, VML), genetic neuromuscular diseases, and premature/accelerated senescence (e.g., sarcopenia) achieving significant results through rehabilitative interventions (Greising, Dearth, and Corona 2016, Roche 2022, Lo et al. 2020). When compared to injecting/implanting myogenic cells into host muscles, implanting actual or engineered muscle tissue along with myogenic cells is more effective in stimulating donor-cell-derived myogenesis (Roche et al. 2020, Roche, Begam, and Galen 2017, Zhang et al. 2014, Moens et al. 1992). Our laboratory has developed a technique known as minimally-invasive muscle embedding (MIME), which promotes robust and consistent donor-cell-derived myogenesis (Roche et al. 2020, Roche, Begam, and Galen 2017). The standard experimental method of performing MIME involves, passing a hypodermic needle (16 or 18 gauge) through the long axis of the tibialis anterior (TA) muscle of a mouse, threading a thin segment of donor muscle tissue with its myogenic cells through the needle track, sealing the needle wounds with the donor tissue embedded within the host muscle, and assessing donor-cell-derived myogenesis several days or weeks later (Roche et al. 2020, Roche, Begam, and Galen 2017). In essence, our data suggest that if myogenic tissue, whether actual or engineered, can be implanted into host muscle through MIME, it is likely to promote donor-cell-derived myogenesis. Recently, our laboratory has attempted to employ the MIME technique to assess if sheets or tubes made of various materials that support cell culture (e.g., gelatin plus glycerol, corn starch plus glycerol, polyvinyl alcohol, cellulose, collagen sheets, etc.) can be implanted into the TA muscle of previously euthanized mice. Our pilot work suggested that the materials would disintegrate while trying to seed cells in vitro or would be too fragile to be implanted by MIME. Due to promising and relevant results published by others involving the encapsulation of myogenic cells in alginate capsules (Scheffold et al. 2023, Ronzoni et al. 2022), we tested a simple workflow to generate extruded alginate tubes that contained MuSCs/myoblasts. We found that our alginate tubes were resilient enough to handle the MIME technique, and that the tubes were able to promote in vitro myoblast outgrowth many days after preparation and maintenance in culture conditions. Since alginate is bioabsorbable over time in vivo, our simple workflow to generate alginate tubes with myogenic cells and implant them into host muscle with the MIME technique can be leveraged by us and others to assess donor-cell-derived myogenesis in various contexts.

## METHODS

### Generation of alginate tubes

Alginate tubes were generated by making a slurry of sodium alginate powder and distilled water or sodium alginate and cell culture media (Ham’s F-10 with 10% fetal bovine serum and 1% penicillin with streptomycin). The final concentration of alginate in the slurry was 10%. Slurry made with water was used to generate alginate tubes for mechanical testing (stress-strain plot, peak stress tolerance, and stiffness). Slurry made with culture media was used to test the ability of alginate tubes to support the seeding of myogenic cells. The slurry was fed into a 10 ml sterile syringe and extruded by an automatic syringe pump. Extrusion was through a sterile 16-gauge hypodermic needle into a 10% calcium lactate solution. Extrusion was at 2 ml/min and assisted by gravity, such that, the extruded alginate slurry was collected in a 10 ml conical tube containing calcium lactate. Calcium lactate provides Ca2+ ions that help polymerize alginate to form a “bioplastic” (Zhang, Cheng, and Ao 2021).

### Mechanical testing

The alginate tubes were transferred from calcium lactate to distilled water and prepared for mechanical testing. In our hands, we found that phosphate buffered saline (PBS) makes the alginate tubes more fragile. Three segments of alginate tube (for three experimental replicates) were made, and black silk sutures were tied to their ends, approximately 12 mm apart (Fig. 1). These sutures were used to attach the alginate tube segments to a force measurement system (300C-LR Dual Mode, Aurora Scientific, Inc., Aurora, ON, Canada). The actuator of the force measurement system was used to apply strain in increments of 1 mm and stress (force) was plotted. Degree of strain was converted into a percentage of initial length for further analyses.

**Figure 1.**
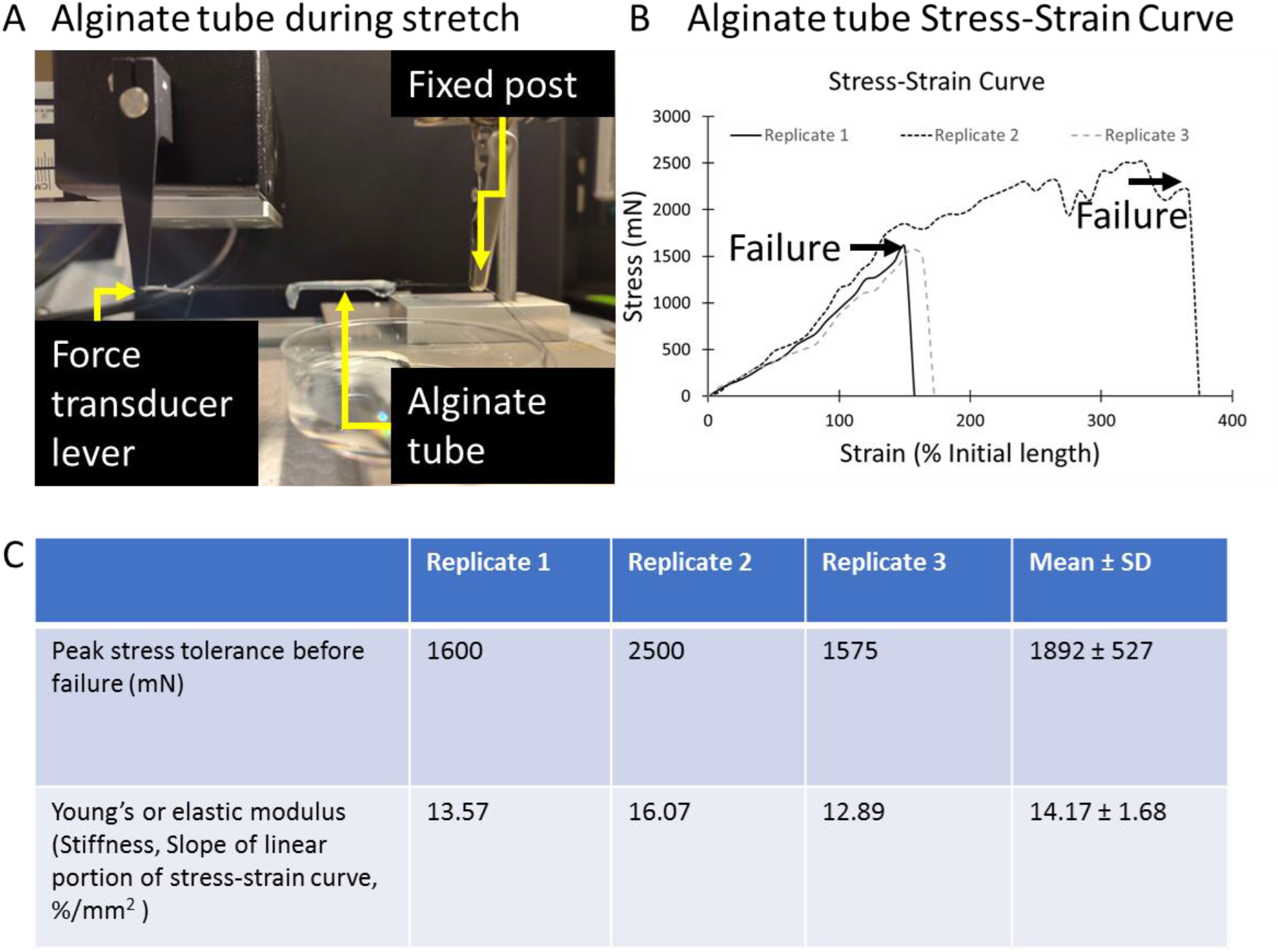
Alginate tube mechanical testing. We have developed the technology to generate cell-encapsulated alginate tubes (A). Briefly, MuSCs harvested from donor muscle tissue are combined with media to give a 10% sodium alginate slurry. The slurry is fed into a syringe and extruded into Ca-lactate through a 16-gauge needle with an automatic syringe pump. Mechanical testing with a force measurement system suggests that the alginate tubes can withstand up to 1500 mN and can undergo strain >150% of its initial length (B, C).

### MuSC isolation

To generate primary myoblast cultures, we followed methods similar to those published by Rando and Blau (Rando and Blau 1994). Briefly, the mouse equivalents of the human quadriceps, hamstrings, gastrocnemius, soleus, TA muscle, pectoralis major, and triceps brachii were harvested from a single ICR/CD1 mouse (female, ~ 6 months, previously euthanized, donated by the Wayne State University Department for Laboratory Animal Resources). Muscles were washed three times with 50 ml sterile PBS in a 50 ml conical tube. The muscles were inverted 10 times in PBS for each wash. Such washing helped remove contaminants, such as hair, and even helped remove some of the adipose tissue that was adhering to muscle. After washing, we manually minced the muscles in the lid of a 60 mm culture dish, with a pair of curved tip scissors, for ~5 min, to generate segments of ~1-2mm. We transferred the minced muscle to a fresh 50 ml tube and performed additional washes with PBS as described above for whole tissue. The additional washing helped remove even more contaminants and adipose tissue (which would float to the top). We then centrifuged (300 RCF, 5 min) the minced tissue with 50 ml PBS in a 50 ml conical tubes and discarded the supernatant liquid. We added an enzyme cocktail (Accumax, STEMCELL Technologies Inc., Cambridge, MA) to the minced tissue at 50% of the volume of minced tissue (e.g., to 5 ml of minced tissue, we added 2.5 ml of enzyme cocktail). The minced tissue plus Accumax was placed on a rocking platform at room temperature for 1 hr per the supplier’s instructions. Following enzyme treatment, we added sterile PBS to the minced tissue and triturated the tissue with a sterile transfer pipette (cut such that the mouth was ~ 5mm wide) to loosen MuSCs. We passed the PBS plus cells through a 70 μm cell strainer into a new 50 ml conical tube. We then inverted the cells with PBS 10 times, and passed it through a 10 μm strainer into a new 50 ml conical tube. We then centrifuged the tube (300 RCF, 5 min) to pellet cells. After discarding the supernatant, we added 10 ml growth media (Ham’s F10, 10% fetal bovine serum, 1% penicillin with streptomycin) to the cell pellet. We divided the cells with growth media into two 5 ml portions – one portion was used to seed cells in three 60 mm dishes to verify that the culture methods were working, and the other portion was made into an alginate slurry for extrusion.

### Maintenance of cells in culture

Cells or tubes containing cells were maintained in culture conditions in a CO2 incubator in 60 mm culture dishes. Cells were in growth media for two days, after which, media was switched to promote fusion (fusion media, Ham’s F10, 2% fetal bovine serum, 1% penicillin with streptomycin). Cells and tubes were incubated in fusion media for six days. The tubes were then implanted into a euthanized animal or were minced (~ 1 cm segments) and reseeded in three 60 mm dishes. The purpose of reseeding cellularized alginate tubes was to assess if the tubes can promote myoblast outgrowth as documented with actual minced muscle (Shahini et al. 2018).

### Implanting alginate tubes into euthanized mice

We performed the MIME technique to implant alginate tube segments into the left TA muscle of mice (ICR/CD1 strain, female, ~ 6 months, previously euthanized, donated by the Wayne State University Department for Laboratory Animal Resources). We implanted one animal with an alginate tube without cells, and implanted another with a cellularized alginate tube. Soon after MIME, we harvested the TA muscles and snap froze them in liquid nitrogen. We performed hematoxylin and eosin staining on the TA muscle cross sections (20 μm thick) to assess the level of cellularization of the alginate tube – hematoxylin staining of nuclei would indicate the presence of cells in the alginate tube.

## RESULTS

**Figure 2.**
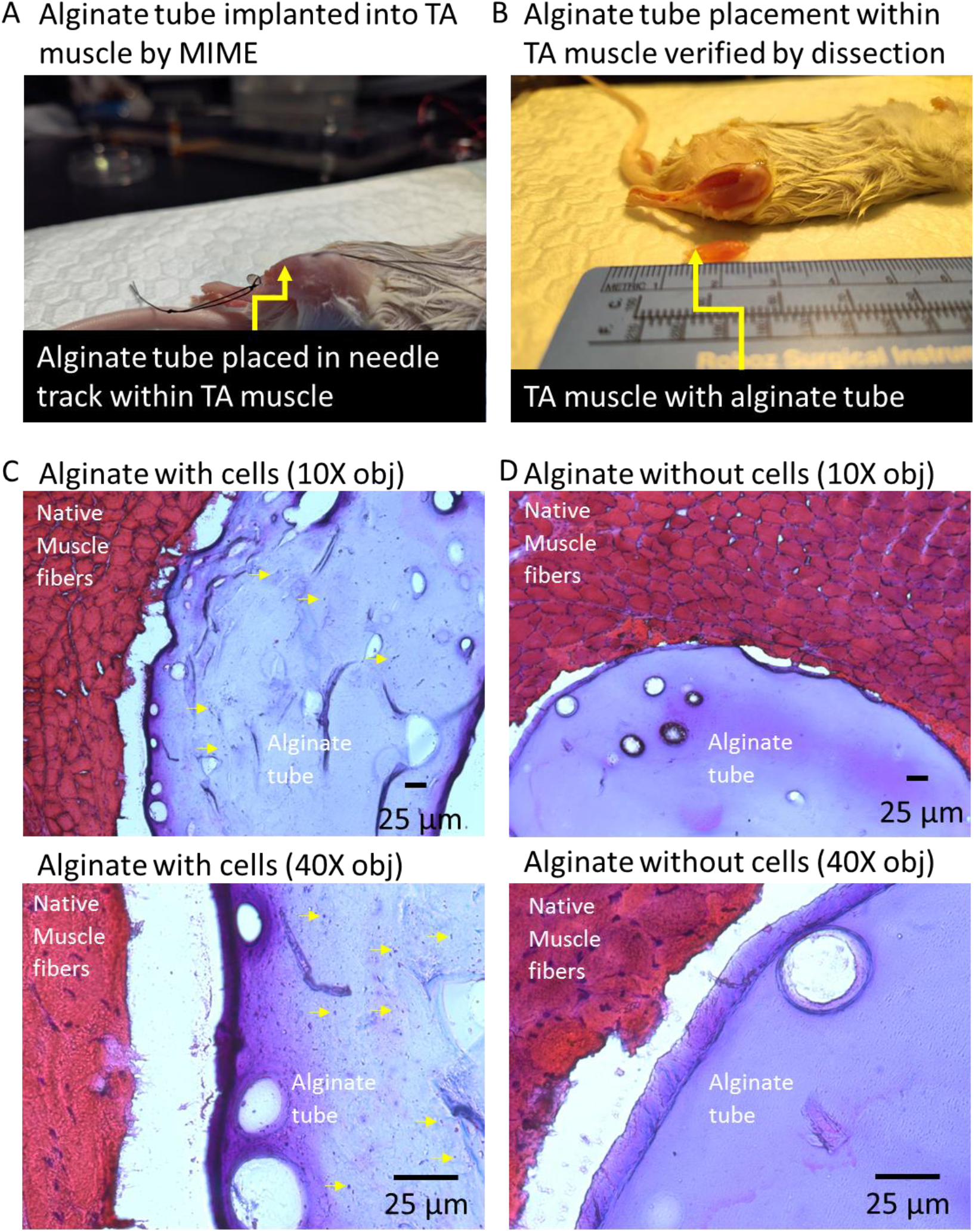
Alginate tubes implanted into the mouse TA muscle by MIME. Alginate (10% sodium alginate) tubes fabricated by a simple extrusion process are amenable to mechanical handling required for implantation into host muscle with the MIME technique (A, B). Frozen, 20 μm cross sections that were stained with hematoxylin and eosin show that the alginate tubes implanted by MIME are within the compartment of the TA muscle (C, D). Cellularized alginate shows evidence of nuclear staining with hematoxylin, whereas alginate that has not been cellularized does not show nuclear staining.

**Figure 3.**
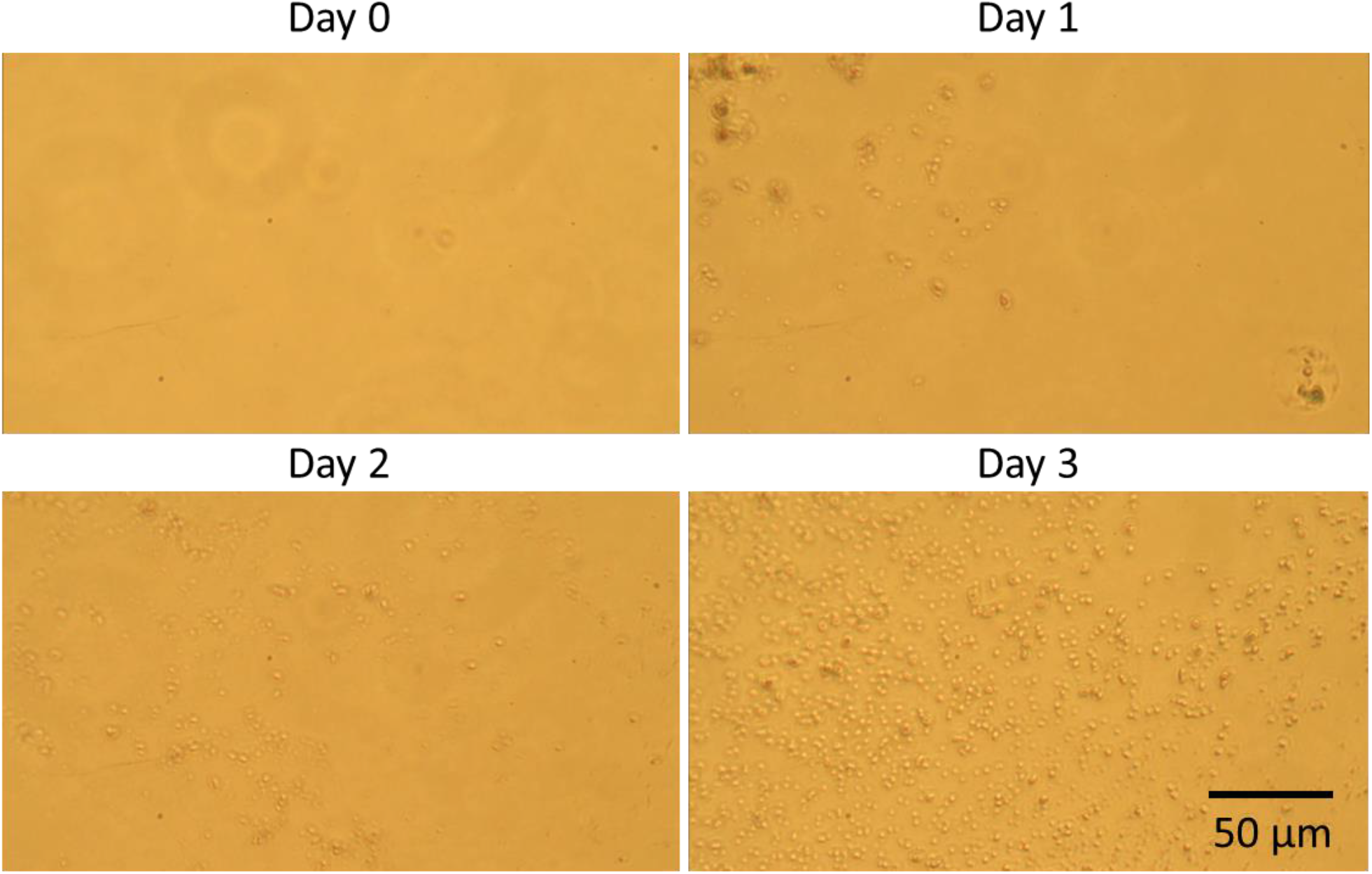
Myoblast outgrowth from minced alginate tubes. Alginate tubes that were cellularized and maintain in cell culture for several days (two days in growth media and six days in differentiation media) were minced and reseeded in 60 mm plates with growth media. The purpose of this experiment was to test if cell-seeded alginate tubes can promote myoblast outgrowth as documented with actual muscle tissue. Our data suggest that over several days after reseeding, minced alginate tubes are able to facilitate myoblast outgrowth as evidenced by increasing confluence over time.

## DISCUSSION

Many advances have been made in the field of regenerative muscle biology. However, the field has not seen a major breakthrough in terms of being able to regrow functional muscle mass in various pathological states (e.g., VML, genetic muscle diseases, sarcopenia). The shift from cell- to tissue-based therapies shows promise in the area of regenerative muscle biology as with other areas of regenerative medicine (Lin et al. 2014, Haraguchi et al. 2012). We have developed a simple workflow to generate cellularized alginate tubes that can be combined with minimally-invasive methods of implantation into host muscle tissue. The simplicity and cost-effectiveness of this workflow could promote global inclusion, diversity, and equity in terms of regenerative muscle biology research. The workflow can also be combined with rehabilitative, pharmacological, and genetic interventions that may augment functional donor-cell-derived myogenesis. The methodology could be made scalable since regenerative treatments can be repeated until sufficient functional muscle mass has been generated/regenerated.

This pilot study is limited in its translational values since experiments were performed in vitro and with previously euthanized mice. Additional rigorous testing is needed to confirm that cellularized alginate tubes can promote the de novo development of donor-cell-derived muscle fibers, which can contribute to contractile force production.

## CONCLUSION

Alginate tubes with MuSC/myoblasts can be generated by a simple extrusion method. The alginate tubes have sufficient mechanical strength to tolerate insertion into a host muscle with minimally-invasive methods. The cellularized alginate tubes demonstrate myogenic potential since they are capable of being maintained in culture conditions for several days, after which they can still facilitate myoblast outgrowth in a dish.

## ACKNOWLEDGMENTS

This work was supported by NIH R03HD091648, Pitch Competition Award from the Alliance for Regenerative Rehabilitation Research and Training (AR3T) from NIH P2CHD086843, and a subcontract from NIH R01AR079884-01 (Peter L. Jones PI) to JAR. The Wayne State Warrior Funder Program and personal funds from JAR also contributed to this work. The authors thank the Wayne State University Doctor of Physical Therapy Program for supporting student research and providing research training through coursework and research experiences. The authors also acknowledge patients enrolled in [Jain Foundation’s dysferlin registry](https://www.jain-foundation.org/patients-clinicians/the-dysferlinopathy-journey/individual-stories/) for sharing their lived experiences on progressive muscle loss.

## Notes

### Competing Interest Statement

The authors have declared no competing interest.

